# Basal activation of astrocytic Nrf2 in neuronal culture media: challenges and implications for neuron-astrocyte modelling

**DOI:** 10.1101/2024.09.18.613665

**Authors:** M.M.O Elsharkasi, B Villani, G Wells, F Kerr

## Abstract

As a gatekeeper of anti-oxidant and anti-inflammatory cell protection, the transcription factor Nrf2 is a promising therapeutic target for several neurodegenerative diseases, leading to the development of Nrf2 activators targeting Keap1-dependent and independent regulatory mechanisms. Astrocytes play a crucial role in regulating neuronal physiology in health and disease, including Nrf2 neuroprotective responses. As neurons require specific conditions for their differentiation and maintenance, most 2D and 3D co-culture systems use medias that are compatible with neuronal differentiation and function, but also ensure astrocyte survival. Few studies, however, assess the molecular adaptations of astrocytes to changes from astrocyte maintenance medias alone, and their subsequent effects on neurons which may represent technical rather than physiological responses.

Our findings show that while Nrf2 can be effectively activated by the Keap1-Nrf2 protein-protein interaction disruptor 18e, and classical Nrf2 activators DMF and CDDO-Me, in human primary cortical astrocyte monocultures, their efficacy is lost in LUHMES neuron-astrocyte co-cultures. Further investigation revealed that the Advanced DMEM/F12-based LUHMES differentiation media maximally induced basal Nrf2 activity in astrocytes alone, compared to astrocyte maintenance media, thus preventing pharmacological activation. Although Neurobasal slightly activated basal Nrf2, this was not significant suggesting that this media has less impact on astrocytic Nrf2 activity relative to Advanced DMEM/F12.

As Nrf2 is a key regulator of oxidative damage and neuroinflammation, modelling these common features of neurodegenerative diseases may be confounded by environments that maximally activate basal Nrf2. Our findings thus suggest caution in media selection for neuron-astrocyte co-culture in disease modelling and therapeutic Nrf2 activator discovery.

## Introduction

Recent studies have investigated the role of astrocytes in neuronal physiology and neurodegeneration using 2 and 3-dimensional (2D and 3D) co-culture systems (Gradisnik and Velnar, 2023; Purushotham and Buskila, 2023). These models strive to closely replicate the complex cellular microenvironment of native tissues (Antunes et al., 2013), and in the context of neuroscience research enable the investigation of biologically relevant interactions between neurons and their surrounding glial cells. Numerous studies have demonstrated that co-culture systems can predict *in vivo* responses (Sreenivasamurthy et al., 2023), and both neuronal maturation (Hedegaard et al., 2020; Kuijlaars et al., 2016) and astrocyte function (Hasel et al., 2017) have been found to be more reflective of their functional morphologies in brain tissue when they are co-cultured *in vitro* compared to their respective cell-specific monocultures. To support neuronal differentiation and maintenance in these complex models, however, investigators use high glucose medias, such as DMEM/F12 or Neurobasal (Luchena et al., 2022; Mor et al., 2021), supplemented with retinoic acid and/or a variety of growth factors and hormones (Bardy et al., 2015). Astrocytes, including primary cells, survive in these conditions (Luchena et al., 2022), but few studies empirically measure their molecular responses to changes from astrocyte maintenance medias and the potential of this alone to impact astrocyte physiology.

Aberrance of homeostatic astrocyte functions, which are key to protecting neurons in stressful conditions (Lane et al., 2018), is an established driver of pathogenesis in several neurodegenerative diseases (Huang et al., 2022). Considerable evidence suggests that Nrf2, the master regulator of antioxidant transcription encoded for by *Nfe2l2*, plays an important role in boosting neuroprotection in such conditions (Osama et al., 2020), but that much of this is due to its’ physiological activation in astrocytes, rather than neurons, with astrocyte-neuronal coupling proving essential for Nrf2-mediated neuronal protection (Bell et al., 2011, 2015; Murakami et al., 2018; Kraft et al., 2004). Stimulation of Nrf2 in astrocytes has been shown to release extracellular glutathione and this is thought to be the key extracellular factor augmenting protection of neurons (Bakshi et al., 2015; Bell et al., 2011; Gupta et al., 2013; Kraft et al., 2004; Shih et al., 2003) *in vitro* (Liddell et al., 2016) and *in vivo* (Bell et al., 2011) in response to Nrf2.

Activation of Nrf2 is therefore a potential therapeutic approach for neuroprotection in Alzheimer’s (Vargas and Johnson, 2009) and Parkinson’s (Gan et al., 2012) diseases, but astrocytes are key to this response. Early clinical studies identified off-target toxicity using electrophilic Nrf2 activators (Lewis et al., 2021; Muñoz et al., 2017) leading to extensive development of novel, selective pharmacology with improved brain penetrance (Sun et al., 2023). Our study has used LUHMES neuron-astrocyte co-cultures to assess the effectiveness of a novel class of Nrf2 activating compounds that disrupt the protein-protein interaction between Keap1 and Nrf2 (Keap1-Nrf2 PPI disruptors) (Georgakopoulos et al., 2017), with the aim of testing their neuroprotective properties in human neuronal systems. We found that Nrf2 can be stimulated by these activators, as well as classical electrophilic compounds, in human primary cortical astrocytes alone, but efficacy is lost in co-culture with neurons. Our further analyses showed that high basal astrocytic Nrf2 activation was induced by maintaining cells in Advanced DMEM/F12-based neuronal differentiation media alone. Other commonly used neuronal culture medias, such as Neurobasal, only mildly stimulated baseline Nrf2 and may be more suitable for use in contact-dependent co-culture systems and astrocyte-conditioned media experiments. This study has important implications not only for researchers investigating Nrf2 therapeutics, but mechanisms by which astrocytes influence neuronal function under oxidative and inflammatory stressors, which activate Nrf2 signalling in health and disease.

## Methods

### Cell culture

All cells used in this study were grown in flasks or plates pre-coated with a 50 μg/mL poly-L-ornithine (Sigma-Aldrich, P3655-10MG) and 1 μg/mL human plasma fibronectin (Sigma-Aldrich, F2006-1MG) coating solution (250 µL/cm^2^), prepared in autoclaved distilled water, for 3 hours (h) at 37°C (95% air and 5% CO_2_). Culture vessels were then washed with distilled water and dried for five minutes before seeding cells. Coating solution was re-used a maximum of two times before discarding. Coated plates were used immediately, but T75 flasks were stored in Phosphate Buffered Saline (PBS) at 4°C for up to one week before use. All complete medias were stored at 4°C for up to one month.

### Primary human astrocytes culture

Primary human cortical astrocytes (ScienCell, SC-1800) were cultured in Complete Astrocyte Media (SC-1801) containing astrocyte growth supplement (5% v/v; #1852), fetal bovine serum (FBS; #0010) (10% v/v) and Penicillin/Streptomycin (Pen/Strep;5% v/v; #0503), all obtained from ScienCell, and maintained at 37°C (95% air and 5% CO_2_). For independent experiments, cells were thawed, at an approximate density of 1 x 10^6^ per cryovial, onto pre-coated T75 flasks. Complete Astrocyte Media was refreshed every two days until 90% confluency, and cells sub-cultured at a split ratio of 1:3 up to passage P3-5 prior to experimental set-up. Cells were detached using 0.05% trypsin-EDTA, at 37°C for one to two minutes, transferring the solution to 5 mL pre-warmed FBS (Fisher Scientific, 11550356), then incubating the flask for a further one to two minutes at 37°C before dislodging remaining cells using Complete Astrocyte Media and combining with those in the FBS. Cells were centrifuged at 1000 x g for eight minutes at 20°C (Labofuge 400R) and either seeded onto pre-coated T75 flasks for maintenance, or onto pre-coated 24-well or 12-well plates, at a density of 15-22.5 x 10^3^ cells/cm^2^, and incubated at 37°C (95% air and 5% CO_2_) for two days prior to treatment of astrocytes alone.

### Lund human mesencephalic (LUHMES) culture

Lund human mesencephalic (LUHMES) cells (ATCC, CRL-2927) were maintained in Proliferation media (Advanced DMEM/F12 (Fisher Scientific, 11540446), N-2 supplement (Fisher Scientific. 12013479) (1:100), Pen/Strep (1:100), L-glutamine (2.5 mM) and Recombinant basic fibroblast factor (40 ng/mL) (PeproTECH, 100-18B-250UG)), in pre-coated T75 flasks (Greiner BioOne, 83.3911.002) at 37°C (95% air and 5% CO_2_). Media was changed every 2-3 days, and cells passaged at 60-70% confluency, and sub-cultured at a ratio of 1:3 until passage P11-12. Cells were dislodged by incubating with pre-warmed 0.05% trypsin-EDTA at 37°C for two to three minutes, before resuspension in complete media (Advanced DMEM/F12, N-2 supplement (1:100), Pen/Strep (1:100) and 2.5 mM L-glutamine), and centrifugation at 500 x *g* for five minutes (Labofuge 400R). Cells were then seeded onto pre-coated T75 flasks for maintenance or astrocyte-containing plates for co-culture experiments as described below.

### Generation of LUHMES neuron-human astrocyte co-cultures

LUHMES and primary human cortical astrocytes were cultured separately in pre-coated T75 flasks as described above, then co-cultures generated using previously published methods (Ratcliffe et al., 2018). On day zero, human astrocytes were seeded (passage 2-5) onto a pre-coated 24-well plate at a density of 2.25 x 10^4^ cells/cm^2^, and LUHMES differentiation media (Advanced DMEM/F12, N-2 supplement (1:100), Pen/Strep (1:100), 2.5 mM L-glutamine, 1 ng/mL tetracycline (Sigma-Aldrich, 64-75-5), 1 mM dibutyryl adenosine 3′, 5′-cyclic monophosphate (dibutyryl cAMP) (Sigma-Aldrich, 60-92-4) and 2 ng/mL GDNF (Pepro Tech, 450-10) was added to LUHMES growing in a separate T75 flask. At day two, LUHMES were re-plated (passage 6-10) onto astrocytes at a density of 15 x 10^4^ cells/cm^2^ using LUHMES differentiation media without GDNF. At day four, media was changed to LUHMES differentiation media without dibutyryl cAMP. Co-cultures were then treated at day six of differentiation.

### Immunostaining

Co-cultures were differentiated, as described above, on pre-coated 13 mm coverslips (Fisher Scientific, 12392128) in a 24-well plate, then fixed in 4% (w/v) paraformaldehyde for 20 minutes at room temperature. Cells were permeabilized using 0.3% (v/v) Triton X100 (Sigma-Aldrich, 126H1030) in 1x PBS (PBST), prior to blocking with 5% (w/v) bovine serum albumin (BSA; Sigma-Aldrich, A9418-5G) in PBST for 20 minutes at room temperature. Cells were incubated with primary antibodies overnight at 4°C in blocking buffer at the following dilutions: anti-ALDH1L1 (1:70; Abcam ab87117), anti-tubulin β-III (1:5000; Biolegend 801213). Alexa Fluor™ 488 anti-mouse and Alexa Fluor™ 594 anti-rabbit (ThermoFisher Scientific #10256302 and #10266352, respectively), secondary antibodies were then added, at 1:5000 dilution in blocking solution, for one hour at room temperature in the dark. Cell nuclei were stained with ProLong^TM^ Diamond Antifade Mountant with DAPI (Thermo Fisher Scientific, P36966). Cells were visualised using a Zeiss LSM 800 Confocal Fluorescence Microscope, and images were captured and processed using ZEN software.

### Nrf2 activating compound treatments

Monocultures of astrocytes or LUHMES neuron-astrocyte co-cultures were treated with 10 μM Keap1-Nrf2 protein-protein interaction (PPI) disruptor, 18e, and 40 μM dimethylfumarate (DMF) or 10 nM CDDO-Me for 28-32 h prior to analysis of NQO1 activity. Concentrations used for all compounds were based on effective activation of Nrf2 in published studies (Bertrand et al., 2015; Georgakopoulos et al., 2017; Jiang et al., 2014) and 0.1% (v/v) DMSO was used as a vehicle control in all experiments. Most treatments were performed in LUHMES differentiation media or Neurobasal medium (Gibco^TM^ Neurobasal (Fisher Scientific, 11570556), 1 X Gibco^TM^ B-27 (Fisher Scientific, 11530536), 20 mM KCl, Pen/Strep (1:100), 2.5 mM L-glutamine) as indicated.

### NAD(P)H dehydrogenase [quinone] 1 activity assay

NAD(P)H Dehydrogenase [Quinone] 1 (NQO1) Activity Assay kit was used to measure the enzymatic activity of the Nrf2 target gene, NQO1, according to the manufacturers’ instructions (Abcam, Ab184867). Following treatment of human primary astrocytes alone or in co-culture with LUHMES neurons, cells were washed with 1x PBS before solubilizing in 150 µLs extraction buffer on ice. Samples were centrifuged at 17,000 x g (Heraeus Instruments, 75005521) for 20 minutes at 4°C. Supernatants were stored at -20°C and retained for further analysis. For normalisation of samples, total protein content was analysed using a Bicinchoninic acid protein assay (Thermo Fisher Scientific, 23227) according to the manufacturers’ instructions for 96-well plates against a serial dilution of BSA standards from 0 to 2 mg/mL, incubation at 37°C for approximately 30 minutes, and absorbance measurement 562 nm (FLUOstar Omega microplate reader).

NQO1 reactions were performed in clear, flat-bottomed 96-well plates, using 2 μg protein per sample in 50 μL extraction buffer and 50 μL reaction buffer. Control reactions were performed using the NQO1 inhibitor, Dicoumarol, to exclude background absorbance. Absorbance was measured at 440 nm, using a FLUOstar Omega microplate reader, every 20-25 seconds over a five-minute period, at room temperature, to establish progression of enzyme kinetics, in addition to endpoint absorbance measurements following the five-minute reaction.

### Nrf2 small interfering RNA (siRNA) knockdown

Dharmacon™ Accell™ siRNA SMARTpools (Horizon Discovery, E-003755-00-0020) were used to knockdown human *Nrf2* (*Nfe2l2*) gene expression according to the manufacturers’ instructions. Human primary cortical astrocytes were cultured at a density of 22.5 x 10^3^ cells/cm^2^ in a pre-coated 12-well plate and incubated at 37°C for 24 h. 100 μM *Nfe2l2* SMARTpool siRNA stock solution was prepared in 1x siRNA Buffer, then diluted to 1 μM in Complete Astrocyte Media (siRNA delivery mix) for treatment of cells for 72 h. A non-targeting siRNA control (1 μM) (Horizon Discovery, B-002000-UB-100) was included as a negative control. All treatments were conducted in duplicate wells in three independent experiments. Efficiency of knockdown was quantified using Real-Time qPCR, as described below.

### Gene expression analysis by quantitative real-time qPCR

Total RNA was extracted from human primary cortical astrocytes, cultured in 12-well plates, using an RNeasy Mini Kit (Qiagen, 74104), according to the supplier’s instructions. RNA was eluted twice in a total of 30 µLs of RNAse-free water and stored at -80°C. RNA concentration was quantified using a NanoDrop spectrophotometer (NanoDrop ND-1000, Thermo Fisher Scientific), with sample A260/280 ratios ranging from 2.01-2.06 and A260/230 ratios from 1.91-2.19, indicating purity, and concentrations from 242-374 ng/µL.

Contaminating genomic DNA was removed from l µg of each RNA sample using DNase I, Amplification Grade (Thermo Fisher Scientific, 18068015), according to the manufacturers’ instructions. Complementary DNA (cDNA) was then synthesised from 0.8 µg of total, DNAse-treated, RNA using an Applied Biosystems™ High-Capacity RNA-to-cDNA™ Kit (10704217, Fisher Scientific), following the supplier’s instructions and incubation with reverse transcriptase in an Applied Biosystems thermocycler (12313653) for 60 minutes at 37°C and five minutes at 95°C to inactivate enzyme. cDNA samples were stored at -20°C until use.

Oligonucleotides specific for amplification of *Nfe2l2* (NM_006164), *GAPDH* (NM_002046) and *RPLP0* (NM_053275) genes were selected from the Primerbank database (https://pga.mgh.harvard.edu/primerbank). Parameters used for selection were as follows: oligo size 18-25 base pair, amplicon size 150-250 base pair, melting temperature (°C) 59-61 and Guanidine/Cytosine ratio 40-60%. Specificity of oligonucleotide sequences were confirmed using the Basic Local Alignment Search Tool (BLAST) of the National Centre for Biotechnology Information (NCBI) BLAST (http://www.ncbi.nlm.nih.gov). Commercial synthesis was conducted by Integrated DNA Technologies (Coralville, USA), and all oligonucleotides were re-suspended in nuclease-free water to prepare 100 pmol/µL stocks. Efficiency of oligonucleotide pairs was assessed by amplifying two-fold serial dilutions of a standard cDNA (**Figure S1**), using the qRT-PCR methods described below. Efficiency values and correlation coefficients (R^2^), calculated from the slope of each standard curve, are depicted in **Table 1**.

**Table 1:**
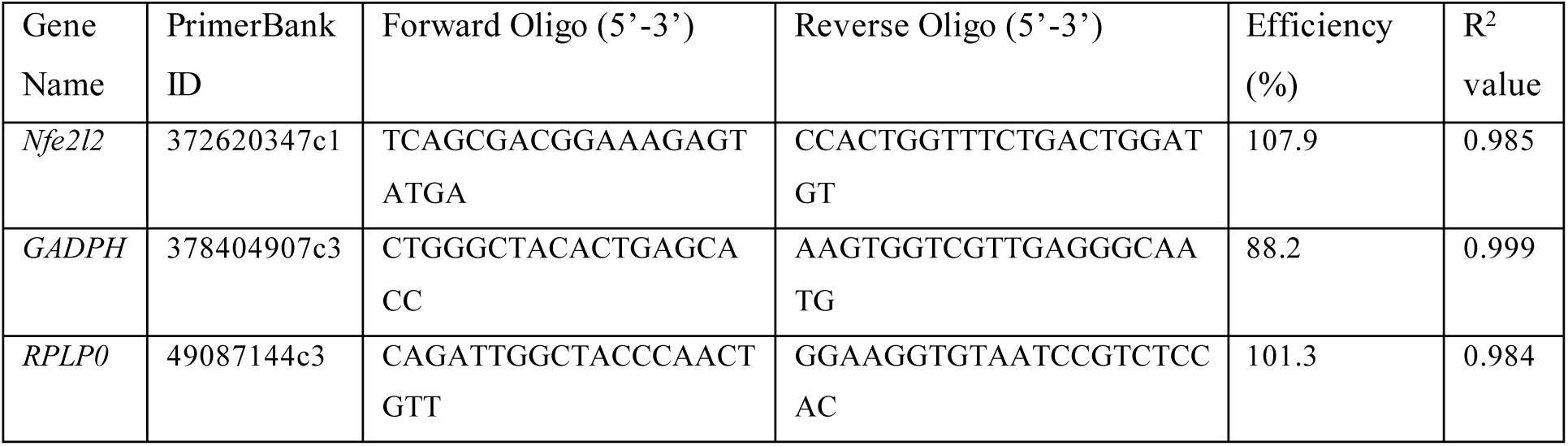
PrimerBank oligonucleotide sequences and efficiency values.

All quantitative RT-PCR steps were performed in a nuclease-free environment in a PCR Workstation (ThermoFisher Scientific, KS12) using UV light to sterilise all equipment and plastics used. qRT-PCR was performed using HOT FIREPol EvaGreen qPCR Mix Plus (Solis BioDyne, 08-24-00001), according to the manufacturer’s recommendations. 20 µLs reactions were prepared directly into white opaque qPCR plates (Primer Design, BW-FAST), comprising 10 ng cDNA, 1 x EvaGreen qPCR master mix, forward and reverse primers (*Nfe2l2* 100 nM, *GAPDH* 250 nM, *RPLP0* 200 nM final concentrations) in nuclease-free water. No template control (NTC) and no reverse transcriptase (RT) controls were included for each gene. The plate was centrifuged at 2000 x g for two minutes at room temperature (Allegra™ X-22R Centrifuge (BECKMAN Coulter, 392187). Three-step amplification reaction was performed using an ABI ViiA 7 Real-Time PCR machine (Thermo Fisher Scientific). The thermal cycling protocol included an initial activation/denaturation at 95°C for 12 minutes, followed by 40 cycles of denaturation at 95°C for 15 seconds, annealing at 60°C for 20 seconds, and extension at 72°C for 20 seconds.

### Data Analysis and Statistics

Three independent experiments were performed for each analysis, with at least two independent wells per experiment. Treatment groups were randomly assigned and samples from individual wells prepared and analysed separately for imaging, protein assays and NQO1 assay. Hence n represents individual wells from at least three independent cell vials or differentiations. This approach follows the NC3Rs recommendations, according to the Reporting *In Vitro* Experiments Responsibly guidelines (The RIVER working group, 2023). Data points from each independent vial/differentiation are denoted by colour and shape in each graph to confirm variance from well to well in each experiment (Lord et al., 2020).

For qRT-PCR analysis, relative expression levels were determined using the arithmetic comparative 2(-Delta Delta C(T)) method (Livak and Schmittgen, 2001). Gene expression was normalised using two reference housekeeping genes, *GAPDH and RPLPO*, and relative expression of the target gene in untreated cells was set to 1.0.

Statistical analyses were conducted using GraphPad Prism 8 statistical software. Normality tests, including the Shapiro-Wilk and D’Agostino-Pearson tests, were performed. Data are expressed as means ± S.E.M and were analysed using two-tailed unpaired t-tests or one-way ANOVA followed by Tukey’s *post-hoc* comparison test for normally distributed data. For data that did not exhibit normal distribution, the Kruskal-Wallis test was conducted, followed by Dunn’s *post-hoc* comparison test. Values were considered statistically significant at *P* < 0.05. Effect size analysis was conducted using the R programming language (R Core Team, 2023; https://www.R-project.org/). The *effsize* package (Torchiano, 2020) was utilised to calculate the Cohen’s d effect size.

## Results

### Pharmacological activation of Nrf2 is effective in astrocytes alone, but not in co-culture with LUHMES neurons

Nrf2 activity was measured indirectly through analysis of enzymatic activity of NQO1, a direct Nrf2 transcriptional target. To ensure observation of NQO1 activity within a linear range and to establish optimal enzyme/substrate ratios for further analysis, assays were performed over a protein concentration range using cell lysates from untreated astrocytes as a source of NQO1 enzyme (**Figure 1A**). The assay was performed using 1, 2 and 4 µg total protein concentrations, with 4 µg being the maximum amount recommended for use with this assay. A dose-dependent increase in NQO1 activity was observed with increasing protein concentration, where 2 and 4 µg protein were significantly increased in comparison to 1 µg (*P* = 0.0066 and 0.0031, respectively) and 2 µg was significantly lower than 4 µg (*P* = 0.0084). Hence, NQO1 activity was detected within the linear range using these methods and 2 µg of total protein per assay was optimal for detection of both increases and decreases in Nrf2 activity.

**Figure 1:**
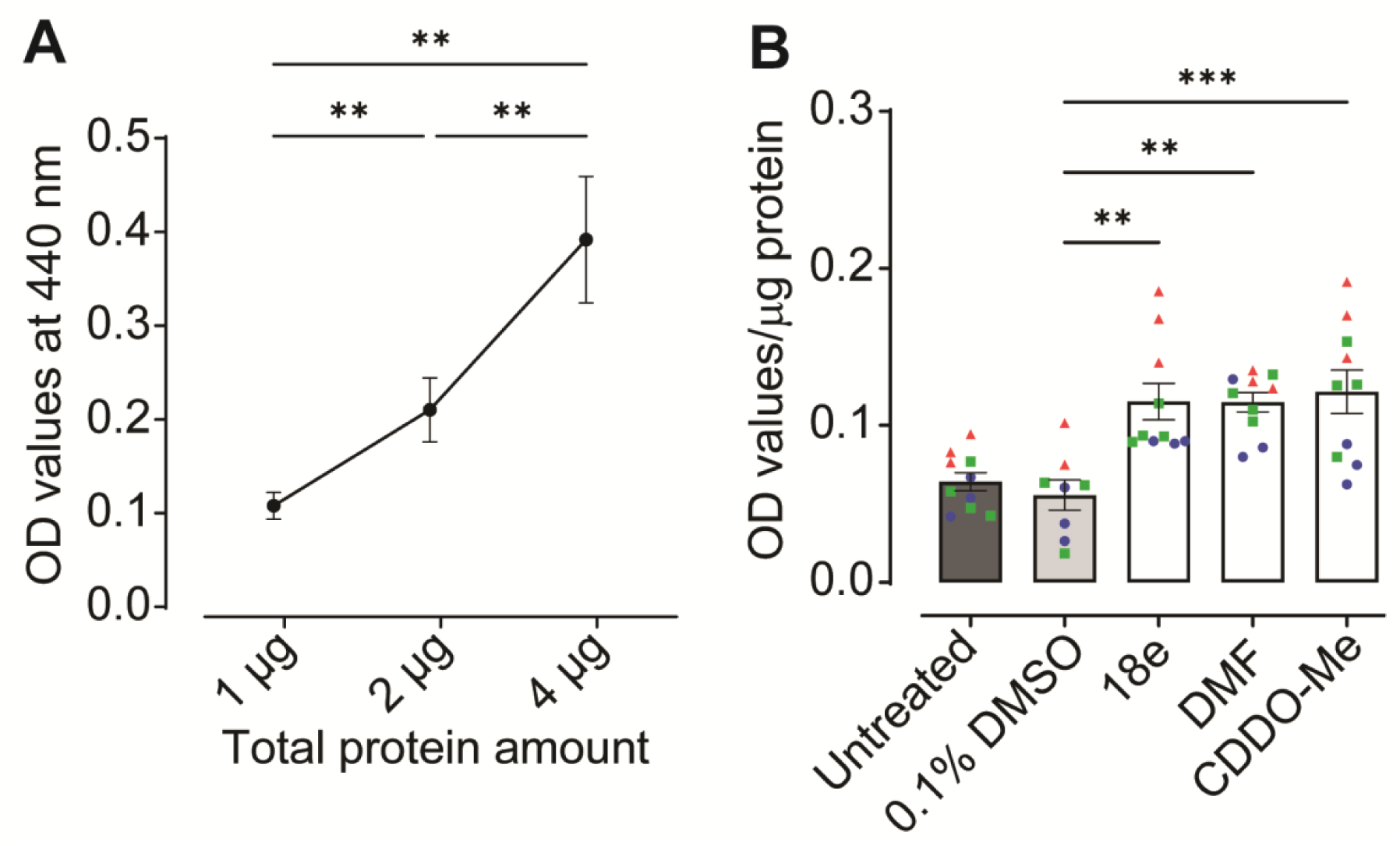
Confirming efficacy of Nrf2 activators in human astrocyte monocultures. **(A)** Activity of NQO1, as measured by simultaneous reduction of its’ substrate menadione and water-soluble tetrazolium salt-1, was used as a read-out of Nrf2 activity. Optimum total protein concentration to measure NQO1 activity in the linear phase was determined using samples from untreated human primary cortical astrocytes. Baseline absorbance, independent from NQO1 activity, was measured in the presence of the NQO1 inhibitor dicoumarol, and OD values subtracted from all data-points presented (n=11, three independent experiments). **(B)** Human astrocytes were treated with pharmacological Nrf2 activators 18e (10 µM), DMF (40 µM) and CDDO-Me (10 nM) for 30-32 h, in ScienCell human astrocyte media, as well as 0.1% DMSO vehicle control (n=8-10, three independent experiments), and Nrf2 activation determined via NQO1 assay using 2 µg total protein per sample. All data are presented as means ± SEM. Statistical analysis was performed using one-way ANOVA followed by Tukey’s multiple comparisons test. **P < 0.01 and ***P < 0.001. Values derived from each independent vial of cells are denoted by colour and shape.

To determine efficacy of a variety of Nrf2 activators, for the purpose of investigating the effects of astrocytic Nrf2 activation on neuronal protection (Bell et al., 2011), primary human cortical astrocytes were first treated with Keap1-Nrf2 PPI disruptor, 18e, and classical electrophilic Nrf2 activators, DMF and CDDO-Me, and NQO1 activity measured (**Figure 1B**). All activators significantly increased NQO1 activity compared to 0.1% DMSO vehicle control (*P* = 0.0017, 0.0019 and 0.0005, 18e, DMF and CDDO-Me respectively; **Figure 1B**). These findings confirm astrocytic activation of Nrf2 in response to electrophilic Nrf2 activators and 18e in human primary cortical astrocytes.

Next, we assessed whether this astrocytic Nrf2 activation was retained when co-cultured in direct contact with LUHMES neurons (**Figure 2**), using methods as described by Ratcliffe et al. (2018). Neuronal differentiation within the co-culture was confirmed by positive immunostaining for the neuronal-specific marker β-III tubulin and human astrocytes were identified using the astrocyte-specific marker aldehyde dehydrogenase 1 family member L1 (ALDH1L1) (**Figure 2A**, I-IV). Specificity of staining was confirmed by lack of non-specific immunofluorescence using secondary antibodies alone or in the absence of primary or secondary antibodies (**Figure 2A**, V-VI). However, activation of Nrf2, as assessed by NQO1 activity, was not observed in LUMHES neuron-astrocyte co-cultures following treatment with either Keap1-Nrf2 PPI disruptors or classical electrophilic activators for up to 32 h (**Figure 2B**). Indeed, high baseline NQO1 activity, with absorbance values >1.0, were observed in untreated co-cultures maintained in LUHMES differentiation media (**Figure 2B**, untreated control). This trended, although not significantly, towards reduced activity, or remained unchanged from DMSO controls, following treatment with all Nrf2 activators. This suggests that Nrf2 is maximally activated in the LUHMES neuron-astrocyte co-cultures, and further activation of the pathway may lead to negative feedback regulation (Occhiuto et al., 2023).

**Figure 2:**
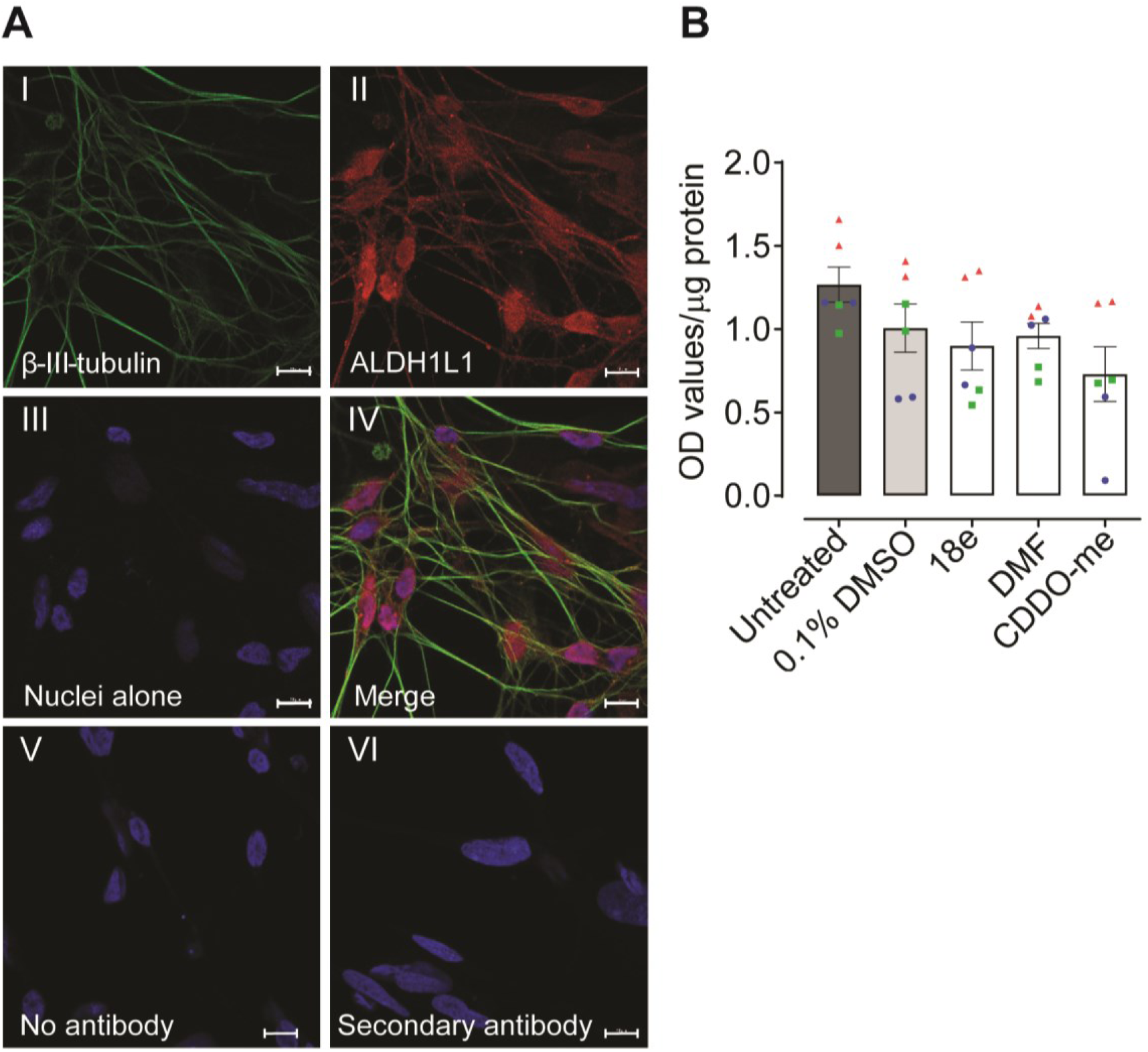
Measuring efficacy of pharmacological Nrf2 activators in LUHMES neuron - astrocyte co-cultures. **(A)** LUHMES neurons were seeded onto a layer of primary human astrocytes at day two of the co-culture protocol, then fixed and immunostained at day six post-differentiation for β-III tubulin (green) and the astrocyte-specific marker ALDH1L1 (red). Nuclei were labelled by DNA staining with DAPI (blue). (I) Neurons alone (II) Human astrocytes alone. (II) Nuclei alone. (IV) Merged image of A-C. (V) No antibody, negative control. (VI) Cells immunostained with goat anti-mouse 488 and goat anti-rabbit 594 secondary antibodies alone. Images were captured using a ZEISS LSM 800 confocal microscope with 63x oil lens. Scale bars, 10 µm **(B)** Effects of Nrf2 activators on NQO1 activity in LUHMES neuron -astrocyte co-cultures maintained in LUHMES differentiation media, following treatment with 18e (10 µM), DMF (40 µM), CDDO-Me (10 nM), and 0.1% DMSO vehicle control, for 25-32 h. (n = 6, three independent experiments). Values derived from each independent vial of cells are denoted by colour and shape.

### Advanced DMEM/F12-based neuronal differentiation media activates NQO1 in human astrocytes through Nrf2-dependent mechanisms

Astrocytes are likely the Nrf2-responsive cell type in the co-culture model (Bell et al., 2011) and thus direct effects of media exposure were further explored using human primary cortical astrocyte monocultures. Given that astrocytes in the co-culture system are maintained in Advanced DMEM/F12-based LUHMES differentiation media, rather than complete astrocyte media, it was hypothesised that a component of this media either activates Nrf2 or induces stress responses in human astrocytes that lead to compensatory increases in baseline Nrf2 activity. We therefore aimed to determine whether LUHMES differentiation media could activate Nrf2 in human primary cortical astrocytes alone (**Figure 3A**&**B**). Astrocyte monocultures were treated with LUHMES differentiation media (untreated control) for 72 h, resulting in an increase in NQO1 activity in comparison with complete astrocyte media, as observed by an increasing rate of enzymatic reaction (**Figure 3A**) and overall activity at the end-point of five-minute reactions (**Figure 3B**; *P* =0.0002 and *P* <0.0001, comparing astrocytes treated with LUHMES Advanced DMEM/F12 differentiation media, alone or in the presence of non-targeting siRNA controls, respectively, to those maintained in astrocyte media).

**Figure 3:**
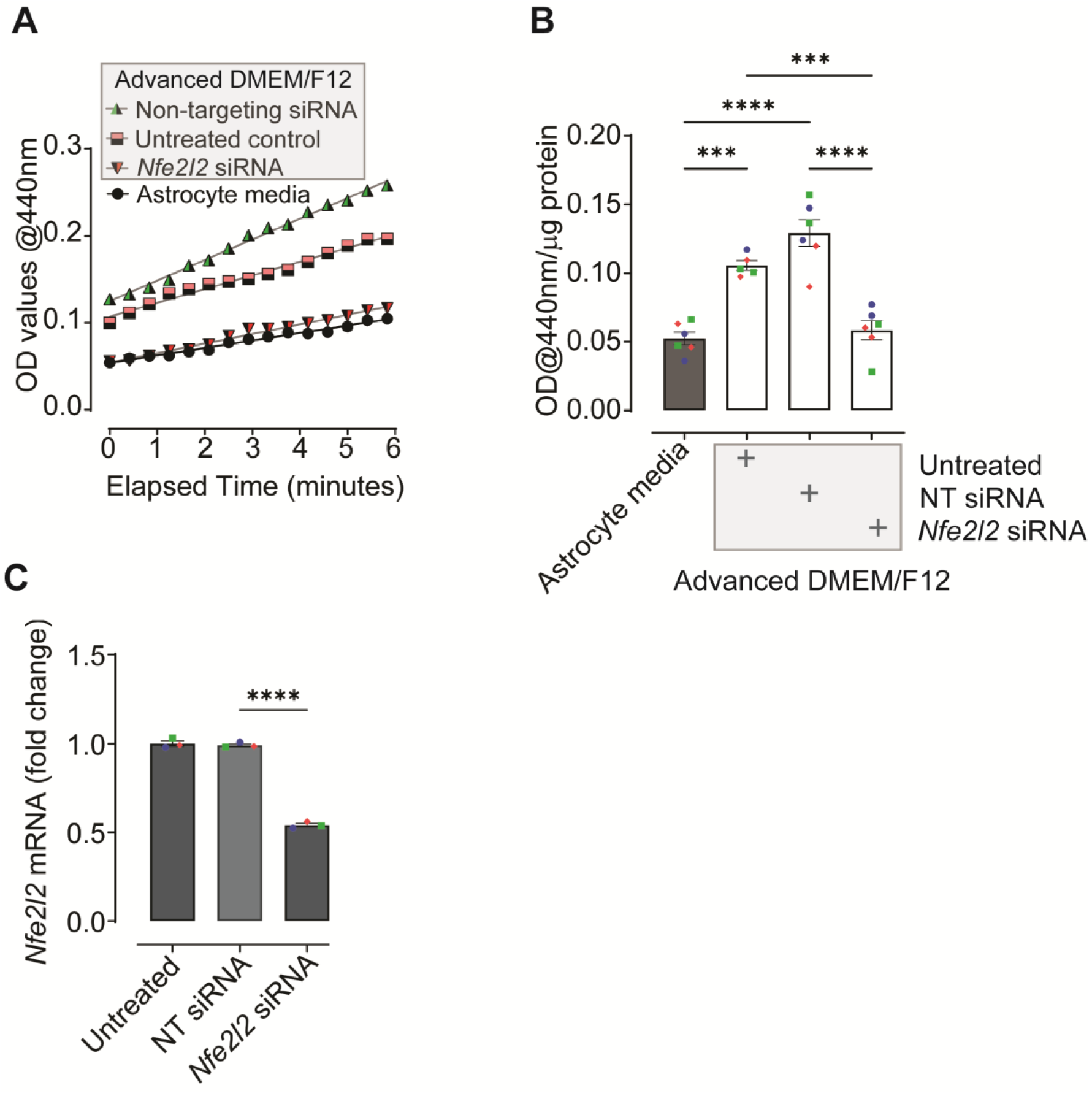
Measuring effects of Advanced DMEM/F12-based LUHMES differentiation media on Nrf2 activity in human primary cortical astrocyte monocultures. **(A)** Kinetic analysis of NQO1 activity. Human cortical astrocytes were treated with LUHMES differentiation media (Advanced DMEM/F12), either alone (untreated control), or in the presence of non-targeting (NT) siRNA or Nfe2I2 siRNAs, and compared to cells maintained in complete astrocyte media. Data are presented as the mean absorbance at 440 nm from three independent experiments, each performed in duplicate. **(B)** NQO1 activity in human primary cortical astrocytes following five-minute reactions. Data are presented as mean absorbance at 440 nm/µg total protein ± SEM (n=6, three independent experiments). **(C)** Nfe2l2 mRNA expression in response to siRNA SMARTpools. Human cortical astrocytes were treated with non-targeting siRNA or Nfe2I2 siRNAs and compared to untreated controls, all prepared in complete astrocyte media. Nfe2l2 gene expression was normalized to the geometric mean of GAPDH and RPLP0 housekeepers, and data are presented as mean fold change ± SEM. (n=3, performed in triplicate, three independent experiments). Analysis performed using one-way ANOVA followed by Tukey’s multiple comparisons test. *P < 0.05, **P < 0.01, ***P < 0.001, and ****P < 0.0001. Values derived from each independent vial of cells are denoted by colour and shape.

Activation of astrocytic NQO1 by LUHMES differentiation media was prevented in the presence of *Nrf2* siRNA SMARTpools (**Figure 3B**; *P* = 0.0008 and *P* <0.0001, Nrf2 siRNA vs Advanced DMEM/F12-based differentiation media alone (untreated control) and in the presence of non-targeting siRNA, respectively), with no significant difference observed between cells cultured in astrocyte media in comparison with cells treated with Advanced DMEM/F12 differentiation media in the presence of Nrf2 siRNAs (*P* = 0.9186). Effectiveness of Nrf2 knockdown using siRNA SMARTpools was confirmed by qRT-PCR for *NFE2L2* mRNA (**Figure 3C**; *P* <0.0001 Nrf2 siRNA-treated vs non-targeting siRNA control), and specificity of effect on NQO1 activity confirmed by observation of no significant differences between non-targeting siRNAs and astrocytes treated with Advanced DMEM/F12 differentiation media alone (**Figure 3B**; *P* = 0.1168). As NQO1 is an indirect measure of Nrf2 activity, our findings therefore verify that induction of this enzyme in astrocytes following treatment with Advanced DMEM/F12-based neuronal differentiation media are indeed due to activation of Nrf2.

### Tetracycline and glucose components of Advanced DMEM/F12-based LUHMES differentiation media have no effect on Nrf2 activity in astrocytes

We next explored potential components of the LUHMES differentiation media, that differed in composition to complete astrocyte media, that could activate Nrf2 in astrocytes. Minocycline, a long-acting second-generation tetracycline analogue, has been previously reported to increase Nrf2 activity in mice (Shahzad et al., 2016). As tetracycline is a key component of Advanced DMEM/F12 LUHMES differentiation media we therefore explored its’effects on Nrf2 activity in human primary cortical astrocytes (**Figure 4**). Cells were treated for 32 h with LUHMES differentiation media with or without 10 µg/mL tetracycline and compared to cells maintained in complete astrocyte media. Under both conditions, NQO1 activity increased, as observed by increasing rates of enzymatic reaction (**Figure 4A**) and overall activity at the end of five-minute reactions, irrespective of the presence of tetracycline (**Figure 4B**; *P* = 0.0101 and *P* = 0.022, comparing Advanced DMEM/F12 differentiation media (+) and (–) tetracycline to astrocyte media control, respectively).

**Figure 4:**
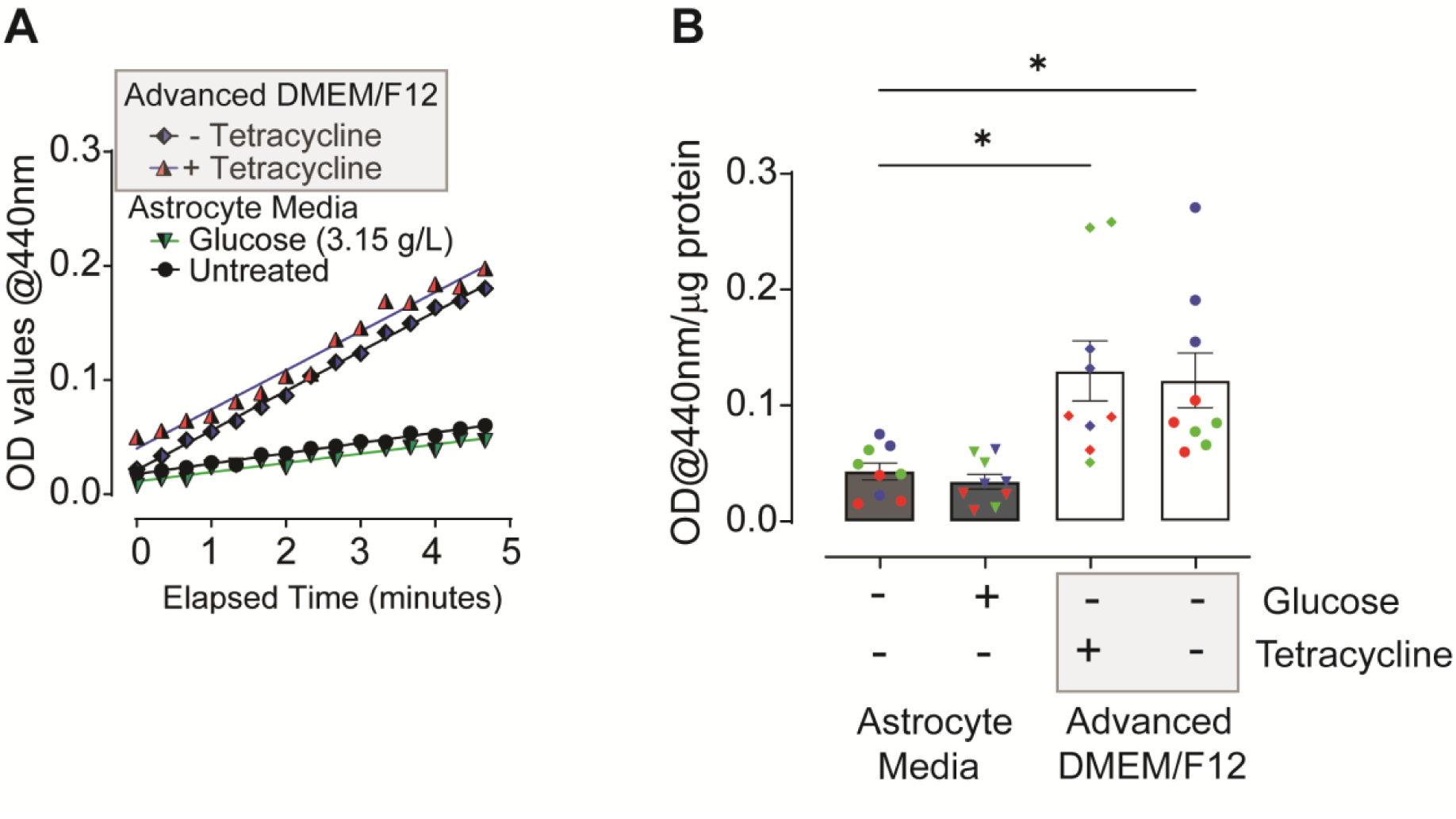
Investigating the effects of Advanced DMEM/F12 LUHMES differentiation media components, glucose and tetracycline, on Nrf2 activity in astrocytes. **(A)** Kinetic analysis of NQO1 activity. Human cortical astrocytes were treated with Advanced DMEM/F12 LUHMES differentiation media, with or without tetracycline (10 µg/mL) or astrocyte media containing 3.15 g/L glucose for 32 h and compared to untreated control cells maintained in complete astrocyte media at 1 g/L glucose. Data are presented as the mean absorbance at 440 nm of three independent experiments, each performed in triplicate. **(B)** End-point analysis of NQO1 activity, after 5 minutes reaction. Data are presented as mean absorbance at 440 nm/µg total protein ± SEM (n=9, three independent experiments). Analysis was performed using one-way ANOVA followed by Tukey’s multiple comparisons test. *P < 0.05. Values derived from each independent vial of cells are denoted by colour and shape.

High glucose has also been shown to activate Nrf2 (Takeda et al., 2021) in human MCF-7 breast cancer cells and to induce changes in the redox regulation of Nrf2 in rat Müller retinal cells (Albert-Garay et al., 2022). As glucose concentrations in LUHMES Advanced DMEM/F12 differentiation media are much higher than those included in complete astrocyte media (3.15 g/L vs 1 g/L), we also investigated whether this could impact on Nrf2 activity in astrocytes. In parallel with the tetracycline studies, human primary cortical astrocytes were therefore maintained in complete astrocyte media either under standard conditions or that supplemented with high glucose (to 3.15 g/L) and NQO1 activity determined (**Figure 4**). Under high glucose (Glucose), no effect on NQO1 activity was observed compared to astrocytes cultured under standard conditions (untreated), based on comparable rates of enzymatic reaction (**Figure 4A**) and overall activity at the end of five-minute reactions (**Figure 4B**; *P* = 0.9858). These findings suggest that neither tetracycline nor glucose influenced Nrf2 activity in human astrocytes.

### Neurobasal media has no significant effect on astrocytic Nrf2 activity

Astrocyte-conditioned media is known to secrete various growth and soluble factors that support neuronal function and offer neuroprotection under physiological conditions and following injury (De Simone et al., 2017; Mahesh et al., 2006; Zhu et al., 2006). However, astrocyte culture media, which commonly contains foetal bovine serum to support growth, is not optimal for differentiation and support of post-mitotic neurons (Scholz et al., 2011). Hence, to explore the effects of astrocyte secretory factors on neuronal cells, astrocytes are commonly conditioned in serum-free media, such as Neurobasal medium, for investigation of non-contact dependent effects on neuronal cultures (Boehler et al., 2007). We therefore assessed the effects of this widely used media (Hagimura et al., 2004; Lopez-Lozano et al., 2022; Scortegagna et al., 1998; Zhang et al., 2022) on basal astrocytic Nrf2 activity and its’ potential to impact on the effectiveness of Nrf2 activators in these cells.

Human primary cortical astrocytes were treated with Neurobasal media for 32 h and compared to cells maintained in complete astrocyte media. Pre-incubation with Neurobasal for 24 h was implemented to allow astrocytes to acclimatise before the treatment period, as is common in astrocyte conditioned media experiments. A slight increase in NQO1 activity in response to neurobasal media was observed compared to complete astrocyte media, as assessed by the rate of the enzymatic reaction (**Figure 5A**) and overall activity at the end of five-minute reactions, but this was not significant (**Figure 5B**; *P* = 0.1950, comparing neurobasal media to complete astrocyte media). Effect size of neurobasal media compared to astrocyte media was 0.645 (95% confidence interval (CI); -0.38 to 1.67). While this is considered a medium effect size, according to Cohen’s criteria (Cohen, 1988), as the CI crossed zero, this confirms findings from the end-point analyses that the effect observed is not statistically significant.

**Figure 5.**
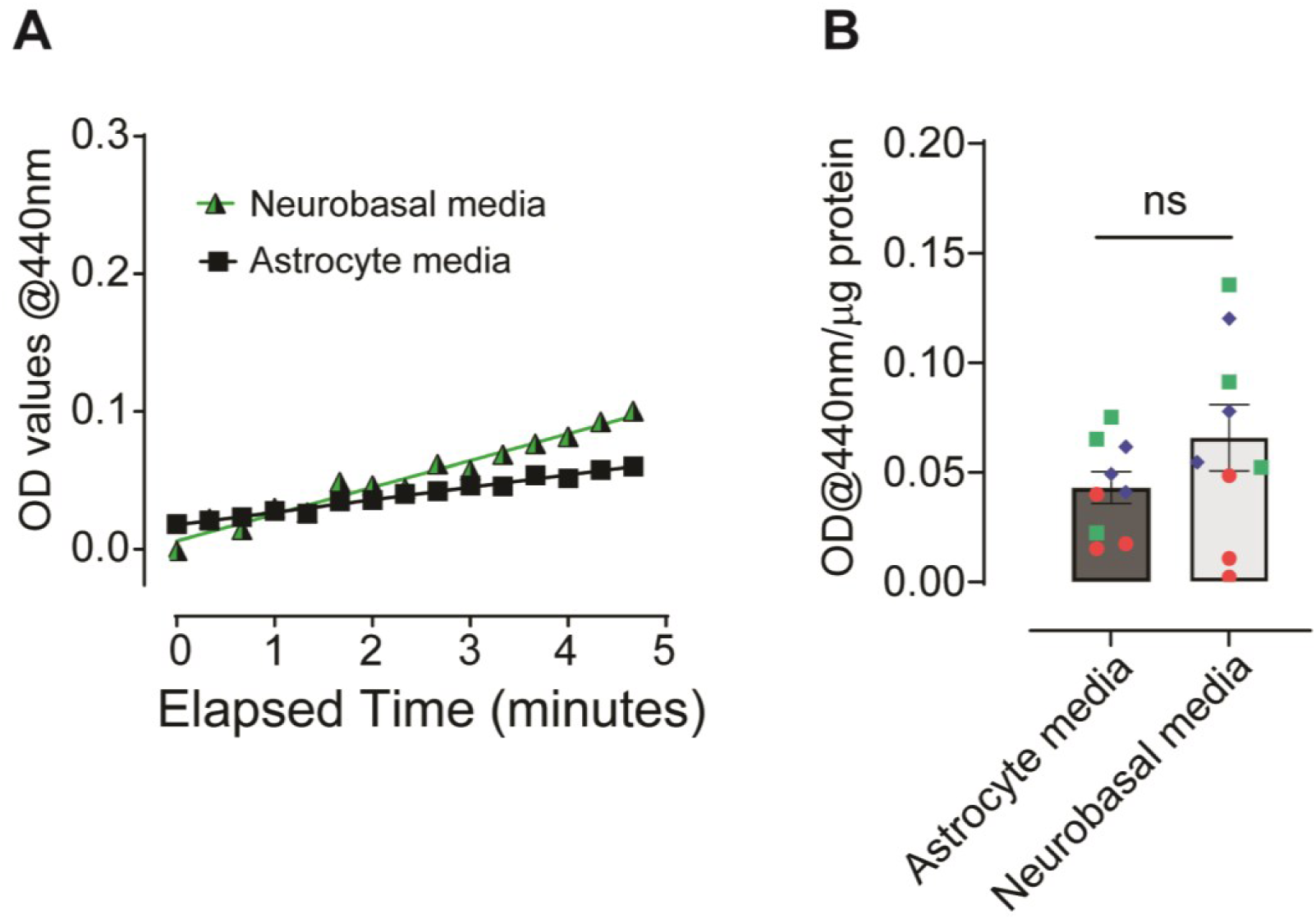
Investigating the effects of Neurobasal medium on Nrf2 activity in human astrocytes. **(A)** Kinetic measurement of NQO1 activity in astrocytes treated with neurobasal media for 32 1. h. **(B)** NQO1 activity after five-minute reactions. Data are presented as mean absorbance/ug total protein ± SEM (n=9, three independent experiments). Analysis in (B) was performed using an unpaired two-tailed t-test and data presented as means ± SEM, *P < 0.05. Values derived from each independent vial of cells are denoted by colour and shape.

## Discussion

Employing 2D and 3D models has emerged as an attractive approach to understanding the overlapping and cell-selective roles of glial cells, such as astrocytes and microglia, in the initiation and progression of several neurodevelopmental (Rehbach et al., 2020; Tomaskovic-Crook et al., 2021) and neurodegenerative conditions (Henstridge et al., 2019; Paumier et al., 2022). Our findings, however, identify the technical complexity of these systems regarding the underlying molecular alterations that could influence functional phenotypes of specific cell types depending on changes in the extracellular environment.

We demonstrate that Nrf2-activating compounds exert robust effects in astrocytes alone but lose efficacy in response to media changes both in monoculture and co-culture with LUHMES neurons. Advanced DMEM/F12-based neuronal differentiation media maximally induced NQO1 activity in astrocytes cultured alone. As enzymatic activity of NQO1 can be affected by a variety of REDOX sensitive pathways that are Nrf2 dependent (Yuhan et al., 2024) or independent (Morrissy et al., 2012), we confirmed that this was due to Nrf2 signalling as media-induced activation was abolished in the presence of *Nfe2l2*-targeting siRNAs.

Neurobasal media, which is used widely for maintaining neurons or preparing astrocyte conditioned media (Boehler et al., 2007; Matafora et al., 2023; Patel and Weaver, 2021; Pozzi et al., 2017), did not significantly induce astrocytic Nrf2 activity, but further studies are required to confirm whether the slight increase in basal activity impacts on efficacy of Nrf2 activators in human astrocytic cells. Of note, our previously published investigations have shown that Nrf2 activity can be robustly induced by the Keap1-Nrf2 PPI disruptor 22h, which is less potent than 18e, DMF and CDDO-Me, in primary mouse neurons maintained in neurobasal (Kerr et al., 2017). These cultures routinely contain approximately 5% astrocytes (Garwood et al., 2011), which contribute the majority of Nrf2 responses in these cells (Al-Mubarak et al., 2021). This suggests that neurobasal does not prevent pharmacological activation of astrocytic Nrf2. Alternatively, this may reflect the more sensitive measurement of direct transcriptional changes of Nrf2 target genes in previous studies (Al-Mubarak et al., 2021; Kerr et al., 2017), which represent an earlier cellular response than alterations in enzymatic activity measured in this investigation.

Our further analyses to establish media components that may induce Nrf2 activation in astrocytes confirmed that this was not due to changes in glucose concentration or addition of tetracycline in Advanced DMEM/F12 neuronal differentiation media. Nrf2 is a master regulator of cellular redox homeostasis and defence against oxidative, inflammatory and xenobiotic stress (Ramsey et al., 2007) and therefore may be indirectly sensitive to environmental stimuli that induce cellular damage. Several investigations suggest that phenol red, used commonly to assess changes in pH in mammalian cell culture, can induce REDOX stress (Morgan et al., 2019) and accumulation of reactive oxygen species (Shvedova et al., 2000). Advanced DMEM/F12, Neurobasal and ScienCell complete astrocyte medias all contain 20-21.5 µM phenol red but differentially induce astrocytic Nrf2 activity in our study, suggesting this is unlikely due to phenol red-induced oxidative damage. Other key supplements, including B-27 in Neurobasal and N2 in DMEM/F12-based medias, contain a complex mix of vitamins, hormones and antioxidants (Sünwoldt et al., 2017) to support neurons, and have been shown to alter cellular metabolism by preventing glycolysis in rat primary neuronal cultures (Sünwoldt et al., 2017). Genetic Nrf2 activation, via Keap1 knockdown, however, increases glucose uptake and mitochondrial NADH levels in mouse primary neurons and astrocytes (Esteras et al., 2023), indicative of increased glycolysis and TCA cycle. This infers that B-27 and N2 are unlikely to activate Nrf2, but their effects on metabolism vary depending on basal media composition, for example losing their effects on glycolysis in rat neuronal cultures when supplementing BrainPhys media (Sünwoldt et al., 2017). As the chemical composition of the ScienCell astrocyte growth supplement is proprietary, but likely forms comparable hormonal and antioxidant support for astrocytes, the baseline effects of these neuronal supporting supplements on astrocytic Nrf2 activity should be considered in future studies.

Nrf2 belongs to the cap’n’collar family of transcription factors and heterodimerises with small Maf proteins to regulate expression of over 250 genes (Osama et al., 2020) containing an Antioxidant Response Element (ARE) (Robledinos-Antón et al., 2019). Physiologically, upregulation of these genes enables cells to maintain REDOX homeostasis, proteostasis and protection from damage (Robledinos-Antón et al., 2019). Canonically, this is tightly regulated under basal conditions by inhibition of Nrf2 through binding to Keap1, which sequesters cytoplasmic Nrf2 and targets it for proteasomal degradation via a Cullin E3-ligase complex (Georgakopoulos et al., 2022; Kerr et al., 2017; Robledinos-Antón et al., 2019). Alternatively, Nrf2 may be independently regulated through phosphorylation by glycogen synthase kinase-3 (GSK-3) and formation of a β-TrCP-CUL1/RBX1 complex which targets it for ubiquitin-proteasome degradation (Jaffry and Wells, 2023; Robledinos-Antón et al., 2019). At the transcriptional level, Nrf2 can be regulated through the presence of a xenobiotic response element, NFkB binding site and ARE in the promoter region of the *Nfe2l2* gene, enabling responses to xenobiotic/inflammatory stimuli and autoregulation to circumvent prolonged stimulation, as well as epigenetic, microRNA and long noncoding RNAs (Robledinos-Antón et al., 2019).

Although our study does not definitively identify specific media components that activate Nrf2, or which induce basal stress responses in astrocytes upon media changes, the mode of action of the pharmacological compounds used may elucidate the mechanisms by which it is regulated in response to Advanced DMEM/F12. Most classical Nrf2 activators, including DMF and CDDO-Me, are electrophilic molecules that modify REDOX-sensitive cysteine residues on Keap1 (Saito et al., 2016) and prevent Nrf2 ubiquitination and degradation (Robledinos-Antón et al., 2019). As these molecules can also modify other REDOX-sensitive proteins they may exert effects independently of Keap1. Thus, development of PPI inhibitors, such as 18e, which interfere with docking of Nrf2 to the Kelch domain of Keap1 (Georgakopoulos et al., 2022; Jiang et al., 2016; Tran et al., 2019) offers increased selectivity for this mechanism of Nrf2 regulation. Our investigation has shown that both electrophilic Nrf2 activators and Keap1-Nrf2 PPI disruptors significantly increased Nrf2 activity in human astrocytes cultured in isolation, but that efficacy is lost in Advanced DMEM/F12 differentiation media both when co-cultured with neurons and alone. This suggests that Advanced DMEM/F12 media activates Nrf2 through disrupting its’ interaction with Keap1, as opposed to other modes of regulation, as compounds that block the Keap1-Nrf2 interaction through distinct, and selective, mechanisms cannot synergistically activate Nrf2.

Media changes that affect Keap1 regulation of Nrf2 specifically may therefore impact on the analysis of other pharmacological and environmental factors that alter Nrf2 activity via this mechanism in astrocytes. This is important because thiol groups on cysteine residues of Keap1 are also subject to modification by reactive oxygen species (Robledinos-Antón et al., 2019), which are a key feature of many neurodegenerative diseases (Houldsworth, 2023). Disease modelling *in vitro* therefore frequently incorporates exposure to oxidative stress, including hydrogen peroxide (Al-Mubarak et al., 2021) and oxygen-glucose deprivation (Baldassarro et al., 2017; Juntunen et al., 2020), or inflammatory cytokines such as TNFα (Hyvärinen et al., 2019; Jayaraman et al., 2021). Studies have shown that neurotoxic effects of oxidative damage may be prevented by conditioned media from either human or mouse astrocytes. Gutbier et al. (2018) have shown this using LUHMES-compatible Advanced DMEM/F12 media to condition astrocytes but, despite demonstrating that this is dependent on glutathione release, to our knowledge few studies have measured basal astrocytic Nrf2 activation in response to neuronal medias. Our findings suggest for the first time that astrocytic Nrf2 activity may require measurement in such investigations to ascertain whether neuronal support is provided due to technical rather than physiological reasons. Alternatively, cellular environments which elevate basal Nrf2 activity may limit the ability of new drugs to activate Nrf2 further, as observed in our investigation, potentially leading to false negative findings where efficacy is masked depending on the cell-type used and pathologies modelled. These potential confounding factors will depend on the mechanisms by which astrocytic Nrf2 is regulated by oxidative or inflammatory stressors, with potential for physiological and pharmacological Nrf2 activation to remain if Keap1-independent. For example, mild oxidative stress following treatment with hydrogen peroxide has been shown to elevate Nrf2 activity in mouse primary astrocytes independently of Keap1, meaning that Keap1 inhibitors can still effectively activate Nrf2 in these conditions (Al-Mubarak et al., 2021). Conversely, molecules which stimulate Nrf2 via Keap1-independent mechanisms, such as lithium and other GSK-3 inhibitors (Kerr et al., 2017, 2018), may still be effective in Advanced DMEM/F12.

Overall, our research suggests that Neurobasal media is likely to be more suitable for investigating the effects of astrocytes on neuronal phenotypes than DMEM/F12-based medias, due to the potential sensitivity of the Nrf2 pathway to changes in the basal extracellular environment. We recommend caution when using common neuronal maintenance medias for astrocyte conditioning or contact-dependent neuronal cultures, to avoid masking Nrf2 responses from pharmacological Keap1-dependent activators or its’physiological upregulation during cellular stress in the study of neurological diseases.

## Notes

### Competing Interest Statement

The authors have declared no competing interest.

### Summary of Updates

Due to an error in analysis, Figure 5C, and accompanying text referring to this figure, has been removed to allow for repetition of the experiments and validation of results. The abstract, introduction and discussion have also been summarised for publication purposes.

## References

Albert-Garay JS, Riesgo-Escovar JR and Salceda R (2022) High glucose concentrations induce oxidative stress by inhibiting Nrf2 expression in rat Müller retinal cells in vitro. Scientific Reports 12(1): 1261.

Al-Mubarak BR, Bell KFS, Chowdhry S, et al. (2021) Non-canonical Keap1-independent activation of Nrf2 in astrocytes by mild oxidative stress. Redox Biology 47: 102158.

Antunes F, Andrade F, Araújo F, et al. (2013) Establishment of a triple co-culture in vitro cell models to study intestinal absorption of peptide drugs. European journal of Pharmaceutics and Biopharmaceutics 83(3): 427–435.

Bakshi R, Zhang H, Logan R, et al. (2015) Neuroprotective effects of urate are mediated by augmenting astrocytic glutathione synthesis and release. Neurobiology of Disease 82: 574.

Baldassarro VA, Marchesini A, Giardino L, et al. (2017) Vulnerability of primary neurons derived from Tg2576 Alzheimer mice to oxygen and glucose deprivation: Role of intraneuronal amyloid-β accumulation and astrocytes. Disease Models and Mechanisms 10(5): 671–678.

Bardy C, Van Den Hurk M, Eames T, et al. (2015) Neuronal medium that supports basic synaptic functions and activity of human neurons in vitro. Proceedings of the National Academy of Sciences of the United States of America 112(20): E2725–E2734.

Bell KF, Al-Mubarak B, Fowler JH, et al. (2011) Mild oxidative stress activates Nrf2 in astrocytes, which contributes to neuroprotective ischemic preconditioning. Proceedings of the National Academy of Sciences of the United States of America 108(1): E1–E2.

Bell KFS, Al-Mubarak B, Martel MA, et al. (2015) Neuronal development is promoted by weakened intrinsic antioxidant defences due to epigenetic repression of Nrf2. Nature Communications 6(1): 7066.

Bertrand HC, Schaap M, Baird L, et al. (2015) Design, Synthesis, and Evaluation of Triazole Derivatives That Induce Nrf2 Dependent Gene Products and Inhibit the Keap1-Nrf2 Protein-Protein Interaction. Journal of Medicinal Chemistry 58(18): 7186–7194.

Boehler, Wheeler BC and Brewer GJ (2007) Added astroglia promote greater synapse density and higher activity in neuronal networks. Neuron Glia Biology 3(2): 127.

Cohen J (1988) Statistical Power Analysis for the Behavioral Sciences Second Edition. Available at: https://www.utstat.toronto.edu/#brunner/oldclass/378f16/readings/CohenPower.pdf (accessed 23 August 2024).

De Simone U, Caloni F, Gribaldo L, et al. (2017) Human Co-culture Model of Neurons and Astrocytes to Test Acute Cytotoxicity of Neurotoxic Compounds. International Journal of Toxicology 36(6): 463–477.

Esteras N, Blacker TS, Zherebtsov EA, et al. (2023) Nrf2 regulates glucose uptake and metabolism in neurons and astrocytes. Redox Biology 62: 102672.

Gan L, Vargas MR, Johnson DA, et al. (2012) Astrocyte-specific overexpression of Nrf2 delays motor pathology and synuclein aggregation throughout the CNS in the alpha-synuclein mutant (A53T) mouse model. Journal of Neuroscience 32(49): 17775–17787.

Garwood CJ, Pooler AM, Atherton J, et al. (2011) Astrocytes are important mediators of Aβ-induced neurotoxicity and tau phosphorylation in primary culture. Cell Death & Disease 2011 *2*:6 2(6): E167–E167.

Georgakopoulos N, Talapatra S, Dikovskaya D, et al. (2022) Phenyl Bis-Sulfonamide Keap1-Nrf2 Protein-Protein Interaction Inhibitors with an Alternative Binding Mode. Journal of Medicinal Chemistry 65(10): 7380–7398.

Georgakopoulos ND, Frison M, Alvarez MS, et al. (2017) Reversible Keap1 inhibitors are preferential pharmacological tools to modulate cellular mitophagy. Scientific Reports 7(1): 10303.

Gradisnik L and Velnar T (2023) Astrocytes in the central nervous system and their functions in health and disease: A review. World Journal of Clinical Cases 11(15): 3385.

Gupta K, Patani R, Baxter P, et al. (2011) Human embryonic stem cell derived astrocytes mediate non-cell-autonomous neuroprotection through endogenous and drug-induced mechanisms. Cell Death & Differentiation 2012 *19*:5 19(5): 779–787.

Gupta K, Chandran S and Hardingham GE (2013) Human stem cell-derived astrocytes and their application to studying Nrf2-mediated neuroprotective pathways and therapeutics in neurodegeneration. British Journal of Clinical Pharmacology 75(4): 907–918.

Gutbier S, Spreng A-S, Delp J, et al. (2018) Prevention of neuronal apoptosis by astrocytes through thiol-mediated stress response modulation and accelerated recovery from proteotoxic stress. Cell Death & Differentiation 25: 2101–2117.

Hagimura N, Tsuzuki K, Iino M, et al. (2004) Predominant expression of GluR2 among the AMPA receptor subunits in neuronal progenitor cells of the rat hippocampus. Brain research. Developmental Brain Research 152(2): 213–223.

Hasel P, Dando O, Jiwaji Z, et al. (2017) Neurons and neuronal activity control gene expression in astrocytes to regulate their development and metabolism. Nature Communications 2017 *8*:*1* 8(1): 1–18.

Hedegaard A, Monzón-Sandoval J, Newey SE, et al. (2020) Pro-maturational Effects of Human iPSC-Derived Cortical Astrocytes upon iPSC-Derived Cortical Neurons. Stem Cell Reports 15(1): 38–51.

Henstridge CM, Hyman BT and Spires-Jones TL (2019) Beyond the neuron-cellular interactions early in Alzheimer disease pathogenesis. Nature Reviews Neuroscience 20(2): 94–108.

Houldsworth A (2023) Role of oxidative stress in neurodegenerative disorders: a review of reactive oxygen species and prevention by antioxidants. Brain Communications 6(1): fcad356.

Huang J, Li C and Shang H (2022) Astrocytes in Neurodegeneration: Inspiration From Genetics. Frontiers in Neuroscience 16: 882316.

Hyvärinen T, Hagman S, Ristola M, et al. (2019) Co-stimulation with IL-1β and TNF-α induces an inflammatory reactive astrocyte phenotype with neurosupportive characteristics in a human pluripotent stem cell model system. Scientific Reports 2019 *9*:*1* 9(1): 1–15.

Jaffry U and Wells G (2023) Small molecule and peptide inhibitors of βTrCP and the βTrCP– NRF2 protein–protein interaction. Biochemical Society Transactions 51(3): 925–936.

Jayaraman A, Htike TT, James R, et al. (2021) TNF-mediated neuroinflammation is linked to neuronal necroptosis in Alzheimer’s disease hippocampus. Acta Neuropathologica Communications 9(1): 1–21.

Jiang ZY, Lu MC, Xu LL, et al. (2014) Discovery of potent Keap1-Nrf2 protein-protein interaction inhibitor based on molecular binding determinants analysis. Journal of Medicinal Chemistry 57(6): 2736–2745.

Jiang ZY, Lu MC and You QD (2016) Discovery and Development of Kelch-like ECH-Associated Protein 1. Nuclear Factor Erythroid 2-Related Factor 2 (KEAP1:NRF2) Protein-Protein Interaction Inhibitors: Achievements, Challenges, and Future Directions. Journal of Medicinal Chemistry 59(24): 10837–10858.

Juntunen M, Hagman S, Moisan A, et al. (2020) In Vitro Oxygen-Glucose Deprivation-Induced Stroke Models with Human Neuroblastoma Cell- and Induced Pluripotent Stem Cell-Derived Neurons. Stem Cells International 2020(1): 8841026.

Kerr F, Sofola-Adesakin O, Ivanov DK, et al. (2017) Direct Keap1-Nrf2 disruption as a potential therapeutic target for Alzheimer’s disease. PLoS Genetics 13(3): e1006593.

Kerr F, Bjedov I and Sofola-Adesakin O (2018) Molecular Mechanisms of Lithium Action: Switching the Light on Multiple Targets for Dementia Using Animal Models. Frontiers in Molecular Neuroscience 11: 405875.

Kraft AD, Johnson DA and Johnson JA (2004) Nuclear factor E2-related factor 2-dependent antioxidant response element activation by tert-butylhydroquinone and sulforaphane occurring preferentially in astrocytes conditions neurons against oxidative insult. Journal of Neuroscience. 24(5): 1101–1112.

Kuijlaars J, Oyelami T, Diels A, et al. (2016) Sustained synchronized neuronal network activity in a human astrocyte co-culture system. Scientific Reports 2016 *6*:*1* 6(1): 1–14.

Lane CA, Hardy J and Schott JM (2018) Alzheimer’s disease. European Journal of Neurology 25(1): 59–70.

Lewis JH, Jadoul M, Block GA, et al. (2021) Effects of Bardoxolone Methyl on Hepatic Enzymes in Patients with Type 2 Diabetes Mellitus and Stage 4 CKD. Clinical and Translational Science 14(1): 299–309.

Liddell JR, Lehtonen S, Duncan C, et al. (2016) Pyrrolidine dithiocarbamate activates the Nrf2 pathway in astrocytes. Journal of Neuroinflammation 13(1): 1–14.

Livak KJ and Schmittgen TD (2001) Analysis of Relative Gene Expression Data Using Real-Time Quantitative PCR and the 2−ΔΔCT Method. Methods 25(4): 402–408.

Lopez-Lozano AP, Arevalo-Niño K, Gutierrez-Puente Y, et al. (2022) SSEA-4 positive dental pulp stem cells from deciduous teeth and their induction to neural precursor cells. Head & Face Medicine 18(1): 9.

Lord SJ, Velle KB, Dyche Mullins R, et al. (2020) SuperPlots: Communicating reproducibility and variability in cell biology. Journal of Cell Biology 219(6).

Luchena C, Zuazo-Ibarra J, Valero J, et al. (2022) A Neuron, Microglia, and Astrocyte Triple Co-culture Model to Study Alzheimer’s Disease. Frontiers in Aging Neuroscience 14: 844534.

Mahesh VB, Dhandapani KM and Brann DW (2006) Role of astrocytes in reproduction and neuroprotection. Molecular and Cellular Endocrinology 246(1–2): 1–9.

Matafora V, Gorb A, Yang F, et al. (2023) Proteomics of the astrocyte secretome reveals changes in their response to soluble oligomeric Aβ. Journal of Neurochemistry 166(2): 346–366.

Mor ME, Harvey A, Familari M, et al. (2021) Neural differentiation medium for human pluripotent stem cells to model physiological glucose levels in human brain. Brain Research Bulletin 173: 141–149.

Morgan A, Babu D, Reiz B, et al. (2019) Caution for the routine use of phenol red – It is more than just a pH indicator. Chemico-Biological Interactions 310: 108739.

Morrissy S, Strom J, Purdom-Dickinson S, et al. (2012) NAD(P)H: Quinone Oxidoreductase 1 Is Induced by Progesterone in Cardiomyocytes. Cardiovascular toxicology 12(2): 108.

Muñoz M, Kulick C, … CK-MS, et al. (2017) Liver injury associated with dimethyl fumarate in multiple sclerosis patients. Multiple Sclerosis Journal. 23(14): 1947–1949.

Murakami S, Miyazaki I and Asanuma M (2018) Neuroprotective effect of fermented papaya preparation by activation of Nrf2 pathway in astrocytes. Nutritional Neuroscience 21(3): 176–184.

Occhiuto CJ, Moerland JA, Leal AS, et al. (2023) The Multi-Faceted Consequences of NRF2 Activation throughout Carcinogenesis. Molecules and Cells 46(3): 176.

Osama A, Zhang J, Yao J, et al. (2020) Nrf2: a dark horse in Alzheimer’s disease treatment. Ageing Research Reviews 64: 101206.

Patel MR and Weaver AM (2021) Astrocyte-derived small extracellular vesicles promote synapse formation via fibulin-2-mediated TGF-β signaling. Cell Reports 34(10): 108829.

Paumier A, Boisseau S, Jacquier-Sarlin M, et al. (2022) Astrocyte–neuron interplay is critical for Alzheimer’s disease pathogenesis and is rescued by TRPA1 channel blockade. Brain 145(1): 388–405.

Pozzi D, Ban J, Iseppon F, et al. (2017) An improved method for growing neurons: Comparison with standard protocols. Journal of Neuroscience Methods 280: 1–10.

Purushotham SS and Buskila Y (2023) Astrocytic modulation of neuronal signalling. Frontiers in Network Physiology 3: 1205544.

Ramsey CP, Glass CA, Montgomery MB, et al. (2007) Expression of Nrf2 in Neurodegenerative Diseases. Journal of Neuropathology and Experimental Neurology 66(1): 75.

Ratcliffe LE, Vázquez Villaseñor I, Jennings L, et al. (2018) Loss of IGF1R in Human Astrocytes Alters Complex I Activity and Support for Neurons. Neuroscience 390: 46–59.

Rehbach K, Fernando MB, Brennand KJ, et al. (2020) Studying Human Neurodevelopment and Diseases Using 3D Brain Organoids. Journal of Neuroscience 40(6): 1186–1193.

Robledinos-Antón N, Fernández-Ginés R, Manda G, et al. (2019) Activators and Inhibitors of NRF2: A Review of Their Potential for Clinical Development. Oxidative Medicine and Cellular Longevity 2019(1): 9372182.

Saito R, Suzuki T, Hiramoto K, et al. (2016) Characterizations of Three Major Cysteine Sensors of Keap1 in Stress Response. Molecular and Cellular Biology 36(2): 271–284.

Scholz D, Pöltl D, Genewsky A, et al. (2011) Rapid, complete and large-scale generation of post-mitotic neurons from the human LUHMES cell line. Journal of Neurochemistry 119(5): 957–971.

Scortegagna M, Chikhale E and Hanbauer I (1998) Lead Exposures Increases Oxidative Stress in Serum Deprived E14 Mesencephalic Cultures. Restorative Neurology and Neuroscience 12(2–3): 95–101.

Shahzad K, Bock F, Al-Dabet MM, et al. (2016) Stabilization of endogenous Nrf2 by minocycline protects against Nlrp3-inflammasome induced diabetic nephropathy. Scientific Reports 6(1): 1–14.

Shih AY, Johnson DA, Wong G, et al. (2003) Coordinate regulation of glutathione biosynthesis and release by Nrf2-expressing glia potently protects neurons from oxidative stress. Journal of Neuroscience 23(8): 3394–3406.

Shvedova AA, Kommineni C, Jeffries BA, et al. (2000) Redox cycling of phenol induces oxidative stress in human epidermal keratinocytes. Journal of Investigative Dermatology 114(2): 354–364.

Sreenivasamurthy S, Laul M, Zhao N, et al. (2023) Current progress of cerebral organoids for modeling Alzheimer’s disease origins and mechanisms. Bioengineering & Translational Medicine 8(2): e10378.

Sun Y, Xu L, Zheng D, et al. (2023) A potent phosphodiester Keap1-Nrf2 protein-protein interaction inhibitor as the efficient treatment of Alzheimer’s disease. Redox Biology 64: 102793.

Sünwoldt J, Bosche B, Meisel A, et al. (2017) Neuronal culture microenvironments determine preferences in bioenergetic pathway use. Frontiers in Molecular Neuroscience 10: 294420.

Takeda T, Ogawa M, Kitamoto J, et al. (2021) High Glucose Enhances Skin Sensitizer-induced NRF2 Activation In Vitro. The Open Toxicology Journal 7(1): 8–17.

The RIVER working group (2023) Reporting In Vitro Experiments Responsibly – the RIVER Recommendations. Epub ahead of print 1 July 2023. DOI: 10.31222/OSF.IO/X6AUT.

Tomaskovic-Crook E, Guerrieri-Cortesi K and Crook JM (2021) Induced pluripotent stem cells for 2D and 3D modelling the biological basis of schizophrenia and screening possible therapeutics. Brain Research Bulletin 175: 48–62.

Torchiano M (2020) effsize: Efficient Effect Size Computation [R package effsize version 0.8.1]. Epub ahead of print 5 October 2020. DOI: 10.5281/zenodo.1480624

Tran KT, Pallesen JS, Solbak SM, et al. (2019) A Comparative Assessment Study of Known Small-Molecule Keap1-Nrf2 Protein-Protein Interaction Inhibitors: Chemical Synthesis, Binding Properties, and Cellular Activity. Journal of Medicinal Chemistry 62(17): 8028– 8052.

Vargas MR and Johnson JA (2009) The Nrf2-ARE cytoprotective pathway in astrocytes. Expert Reviews in Molecular Medicine 11: e17.

Yuhan L, Khaleghi Ghadiri M and Gorji A (2024) Impact of NQO1 dysregulation in CNS disorders. Journal of Translational Medicine 2023 *22*:*1* 22(1): 1–28.

Zhang Y, Sun YY, Xu M, et al. (2022) The Stem Cell Potential of O-2A Lineage Astroglia. Developmental neuroscience 44(6): 487–497.

Zhu ZH, Yang R, Fu X, et al. (2006) Astrocyte-conditioned medium protecting hippocampal neurons in primary cultures against corticosterone-induced damages via PI3-K/Akt signal pathway. Brain Research 1114(1): 1–10.

