## Supplementary information for "Basal activation of astrocytic Nrf2 in neuronal culture media: challenges and implications for neuron-astrocyte modelling"

### Supplementary Figures

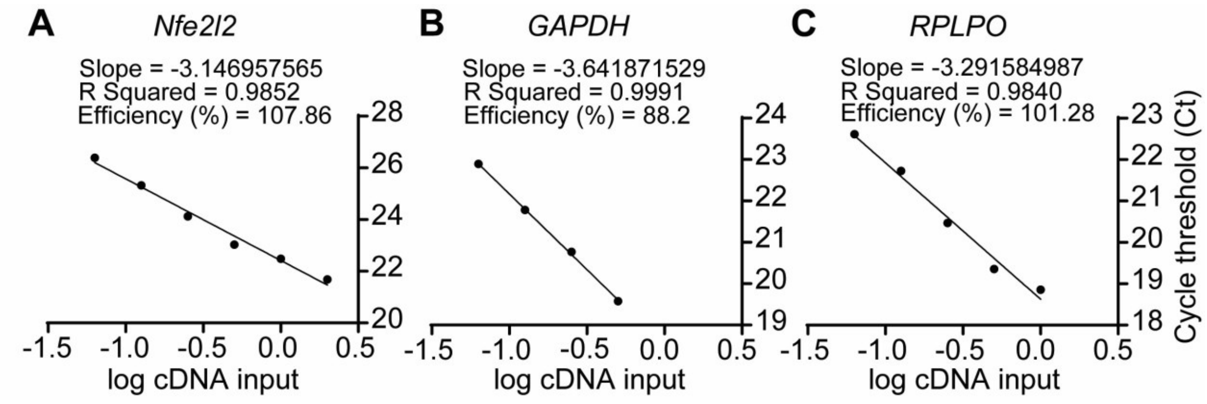


**Figure S1: Standard Curve optimization of A. *Nfe2l2*, B. *GAPDH* and *C. RPLPO* primers.** cDNA from primary human cortical astrocytes was serially diluted, two-fold, over the arbitrary amounts depicted with neat cDNA representing a value of 1.0 (log value 0). qRT-PCR was performed with each primer pair using diluted cDNA and plotted as cycle thresholds (Ct) against log10 cDNA input amount. Slopes were extrapolated from each linear standard curve, and used to calculate efficiency values, using the formula Efficiency = (10e(-1/SLOPE)-1) x 100, and correlation coefficients ($R^2$).
